# Assessing Functional Androgen Receptor Pathway Activity in Response to Radiotherapy Using hK2-targeted PET Imaging

**DOI:** 10.1101/2022.06.23.497290

**Authors:** Claire M Storey, Mohamed Altai, Mesude Bicak, Darren R Veach, Katharina Lückerath, Gabriel Adrian, Michael R McDevitt, Teja Kalidindi, Julie E Park, Ken Herrmann, Diane Abou, Wahed Zedan, Norbert Peekhaus, Robert J Klein, Robert Damoiseaux, Steven M Larson, Hans Lilja, Daniel Thorek, David Ulmert

## Abstract

External beam radiotherapy (EBRT) remains a common treatment for all stages of PCa, but DNA damage induced by EBRT upregulates androgen receptor (AR) pathway activity to promote therapeutic resistance. [^89^Zr]11B6-PET is a novel modality targeting prostate-specific protein human kallikrein 2 (hK2), which is a surrogate biomarker for AR activity. Here, we studied if [^89^Zr]11B6-PET can accurately assess EBRT-induced AR activity. PCa mouse models received EBRT (2-50 Gy) and treatment response was monitored by [^89^Zr]11B6-PET/CT. Radiotracer uptake and expression of AR and AR target genes was quantified in resected tissue. EBRT increased AR pathway activity in LNCaP-AR tumors. EBRT increased prostate-specific [^89^Zr]11B6 uptake and hK2 levels in PCa-bearing mice (Hi*-Myc* x Pb_*KLK2*) with no significant changes in uptake in healthy (Pb_*KLK2*) mice. Thus, [^89^Zr]11B6-PET specifically detects activation of AR pathway activity after EBRT in PCa. Further clinical evaluation of hK2-PET for monitoring EBRT is warranted.

## Introduction

From the maturation of the prostate at adolescence through all stages of prostate adenocarcinoma, the androgen receptor (AR) hormone circuit governs growth, survival and progression of prostate cells^1^. As the central driver of disease progression in prostate cancer (PCa), AR has been the primary target for PCa treatment with the goal of achieving AR signaling inhibition (ARSI) using steroid deprivation or antiandrogens^2^. Alongside ARSI, external beam radiotherapy (EBRT) is an effective treatment option for localized advanced PCa^3^. Inhibition of AR signaling through adjuvant, concurrent, or long-term androgen deprivation treatment increases EBRT efficacy^4,5^. EBRT can directly activate AR and expression of DNA repair genes^6,7^, likely explaining the synergy between ionizing radiation and endocrine therapy.

Resistance to ARSI develops due to aberrant activation of AR signaling from AR overexpression, emergence of AR variants, and intratumor steroidogenesis^8,9^. Continuous AR signaling increases expression of target genes associated with PCa tumor progression and DNA repair to promote radio-resistance^8–11^. Positive feedback loops initiated by DNA repair genes such as PARP-1 and DNA-PKcs further drive resistance by repairing radiation-induced DNA breaks while increasing AR expression^6,7^. These mechanisms enhance the ability of AR-driven PCa cells to accelerate repair of DNA and increase survival after ionizing radiation^6,7^. Furthermore, prostate tumors vary in their AR pathway activity at baseline and after EBRT; analyses of transcriptional signatures of primary PCa biopsies post-EBRT demonstrate varying degrees of AR-pathway activation and heterogeneity in the DNA-damage response between tumors^6^. These variances in radiosensitivity and DNA damage response have been associated with PCa outcome^12^ and response to combinatorial EBRT and ARSI^6,13^.

Non-invasive biomarkers for monitoring DNA damage-induced AR activity would aid in detection of resistance to treatment and provide actionable evidence for drug development and individualized patient treatment options. In the clinical setting, AR activity is currently monitored through the assessment of serum prostate specific antigen (PSA, *KLK3*) levels over time^14^. PSA is the most widely used precision biomarker in oncology and is an extremely sensitive measure of AR-activity through the production and extracellular release of prostate-specific kallikreins^15^. However, measurements of serum kallikreins provide limited information as only a one-millionth fraction of the proteins produced by PCa tissues are released into the blood circulation^16^ and reflect a global average of multiple heterogenic lesions in the metastatic setting with limited correlation to protein production.

DNA damage-induced AR signaling provides an opportunity for monitoring AR pathway activity through downstream target genes. Like PSA, human kallikrein 2 (hK2; *KLK2*) is a prostate gland- and cancer cell-specific trypsin-like serine protease that is tightly governed by the functional status of the AR hormone response circuit. Indeed, EBRT elevates hK2 serum levels in >20% of patients^12^. We previously developed 11B6, an IgG1 antibody with high selectivity and specificity for the active cleavage site of hK2. 11B6 uniquely binds to hK2 directly at the cell surface and avoids interaction with serum kallikreins. When derivatized with medically relevant radionuclides, this platform can be used for radio-immunotheranostics for detection, delineation, and treatment of diverse models of AR-expressing adenocarcinoma^17–19^. Positron emission tomography (PET) with [^89^Zr]11B6 enables quantification of lesion-specific AR-activity^17–19^. Based on our previous experience with the application of [^89^Zr]11B6-PET to monitor disease and observations of hK2 production following irradiation^12,17^, we hypothesize that [^89^Zr]11B6-PET could be used to noninvasively monitor EBRT-induced changes in AR-activity in PCa. Using quantitative imaging and genomic analyses of human xenograft and genetically engineered mouse models of PCa, EBRT-induced AR activity was visualized and correlated to transcriptomic alterations following therapy with near-term implications for PCa treatment paradigms.

## Results

### Changes in AR and AR-driven PCa biomarkers in response to EBRT

PCR analysis of LNCaP-AR tumors treated with 1, 3 or 5 fractions of 2, 5 or 10 Gy EBRT revealed dose-dependent increases in *AR, KLK2*, KLK3 compared to nontreated (NT) controls (**Fig. 1, Tbl. 1**). *FOLH1* expression after EBRT varied and remained unchanged under EBRT (**Fig. 1B, Tbl. 1**). After 3 cycles of EBRT in 22Rv1 xenografts, *AR* gene expression was significantly increased along with *KLK2* and *KLK3*, while there were no significant changes in *FOLH1* expression (**Fig. 1C, Tbl. 1**). The fold change of AR transcription was higher in 22Rv1 than LNCaP-AR tumors, which is likely an effect of lower baseline AR expression in the 22Rv1 model. This outcome corresponds with previously reported findings and provides additional support for the correlation between *KLK2* and *AR* expression when monitoring changes rendered by EBRT^12^.

**Figure 1.**
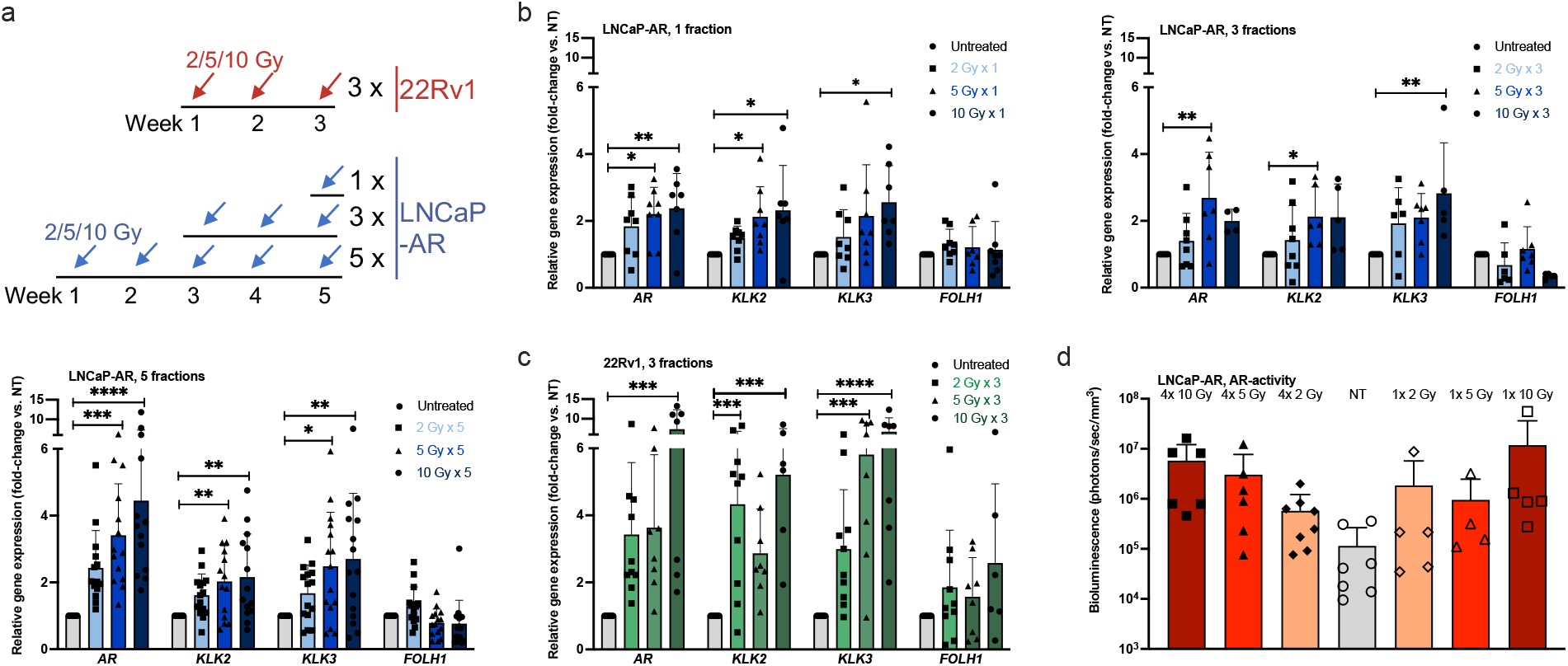
AR activity and gene expression after EBRT in LNCaP-AR and 22Rv1 xenografts. (a) Schematic of EBRT fractionation regimen. (b, c) Gene expression analysis of LNCaP-AR (b) and 22Rv1 (c) xenografts after 1, 3, and 5 fractions of 2, 5 or 10 Gy EBRT revealed upregulation of *AR* and *KLK2*/*KLK3* in a dose-dependent manner. Data were normalized to NT. See Tbl. 1 for mean and p-values. (d) Bioluminescence imaging readout of AR activity in LNCaP-AR xenografts after 1 or 4 fractions of EBRT revealed dose-dependent increase in AR activity independent of fractionation (all p=not significant vs. NT). Mean ± SD and individual values are given; statistical significance was calculated using one-way ANOVA and Dunnett’s test for multiple comparisons.

**Table 1.** Fold-change of AR and AR pathway genes in LNCaP-AR and 22Rv1 tumors after EBRT (vs. controls). Mean ± SD are given; p-values (treatment vs. NT) are shown in parentheses and were calculated using one-way ANOVA and Dunnett’s test for multiple comparisons.

|  | 1 Fraction | 2 Gy | 5 Gy | 10 Gy |
| --- | --- | --- | --- | --- |
| LNCaP-AR | *Ar* | 1.8 ± 0.9<br>(0.1044) | 2.2 ± 0.8<br>*(0.0128) | 2.4 ± 1.0<br>**(0.0057) |
| | *Klk2* | 1.5 ± 0.4<br>(0.4988) | 2.1 ± 0.9<br>*(0.0248) | 2.3 ± 1.3<br>*(0.0102) |
| | *Klk3* | 1.5 ± 0.8<br>(0.6229) | 2.1 ± 1.5<br>(0.0855) | 2.6 ± 1.1<br>*(0.0183) |
| | *Folh1* | 1.3 ± 0.4<br>(0.5323) | 1.2 ± 0.6<br>(0.8127) | 1.1 ± 0.8<br>(0.9315) |
|  | **3 Fractions** | **2 Gy** | **5 Gy** | **10 Gy** |
| | *Ar* | 1.4 ± 0.8<br>(0.6799) | 2.7 ± 1.4<br>**(0.0022) | 2.0 ± 0.4<br>(0.1640) |
| | *Klk2* | 1.4 ± 1.0<br>(0.6169) | 2.1 ± 0.9<br>*(0.0471) | 2.1 ± 1.0<br>(0.0688) |
| | *Klk3* | 1.9 ± 1.1<br>(0.1771) | 2.1 ± 0.7<br>(0.0737) | 2.8 ± 1.5<br>**(0.0052) |
| | *Folh1* | 0.7 ± 0.7<br>(0.4646) | 1.2 ± 0.7<br>(0.8600) | 0.3 ± 0.1<br>(0.0615) |
|  | **5 Fractions** | **2 Gy** | **5 Gy** | **10 Gy** |
| | *Ar* | 2.4 ± 1.1<br>(0.0637) | 3.4 ± 1.5<br>*** (0.0008) | 4.4 ± 2.8<br>**** (<0.0001) |
| | *Klk2* | 1.6 ± 0.6<br>(0.1540) | 2.0 ± 1.1<br>** (0.0079) | 2.2 ± 1.3<br>** (0.0025) |
| | *Klk3* | 1.7 ± 0.8<br>(0.3805) | 2.5 ± 1.6<br>* (0.0104) | 2.7 ± 2.0<br>** (0.0028) |
| | *Folh1* | 1.3 ± 0.6<br>(0.2652) | 0.8 ± 0.4<br>(0.5133) | 0.8 ± 0.7<br>(0.4302) |
| 22Rv1 | **3 Fractions** | **2 Gy** | **5 Gy** | **10 Gy** |
| | *Ar* | 3.4 ± 2.2<br>(0.1296) | 3.6 ± 2.2<br>(0.1180) | 7.3 ± 5.0<br>*** (0.0001) |
| | *Klk2* | 4.3 ± 2.5<br>*** (0.0007) | 2.9 ± 1.3<br>(0.0894) | 5.2 ± 2.3<br>*** (0.0002) |
| | *Klk3* | 3.0 ± 1.8<br>(0.1732) | 5.8 ± 3.2<br>*** (0.0005) | 6.7 ± 3.5<br>**** (<0.0001) |
| | *Folh1* | 1.9 ± 1.7<br>(0.4425) | 1.6 ± 1.2<br>(0.7579) | 2.6 ± 2.4<br>(0.1146) |

Investigating ERBT-induced transcriptomic changes in an unbiased approach, 4,851 DEGs (8.2% of transcriptome gene set) were identified in LNCaP-AR tumors after EBRT (5 × 10 Gy; vs. NT); 2,552 genes were up- and 2,299 were downregulated (**Fig. 2**). Upregulation of AR-regulated genes such as AR signaling co-activator *ETV1* ^20^, *KLK2*, and *KLK3* (log2 fold-change= 10.01, 1.033, 1.882) indicated that AR signaling was increased after EBRT. Interestingly, other AR target genes, including *TMPRSS2* and *FKBP5*, were downregulated following treatment. Of the 144 previously established AR-associated DNA repair genes^6^, 18 were DEGs with 8/18 upregulated (*CHEK1, FANCL, MAD2L1, MBM7, PARP1, RAD18, RAD21, RFC3*)^6,21^. *FOLH1* was also upregulated despite its inverse correlation to AR pathway activity, contrasting qPCR findings. Upregulated *MYC* expression in EBRT-treated tumors supports a role for MYC in AR-driven EBRT responses, and pathway analysis showed that the top DEGs converged on cell cycle and regulation of DNA replication, both of which are closely intertwined with AR through cyclins and changes in protein expression during replication^22,23^, further supporting a role for AR signaling in PCa response to EBRT.

**Figure 2.**
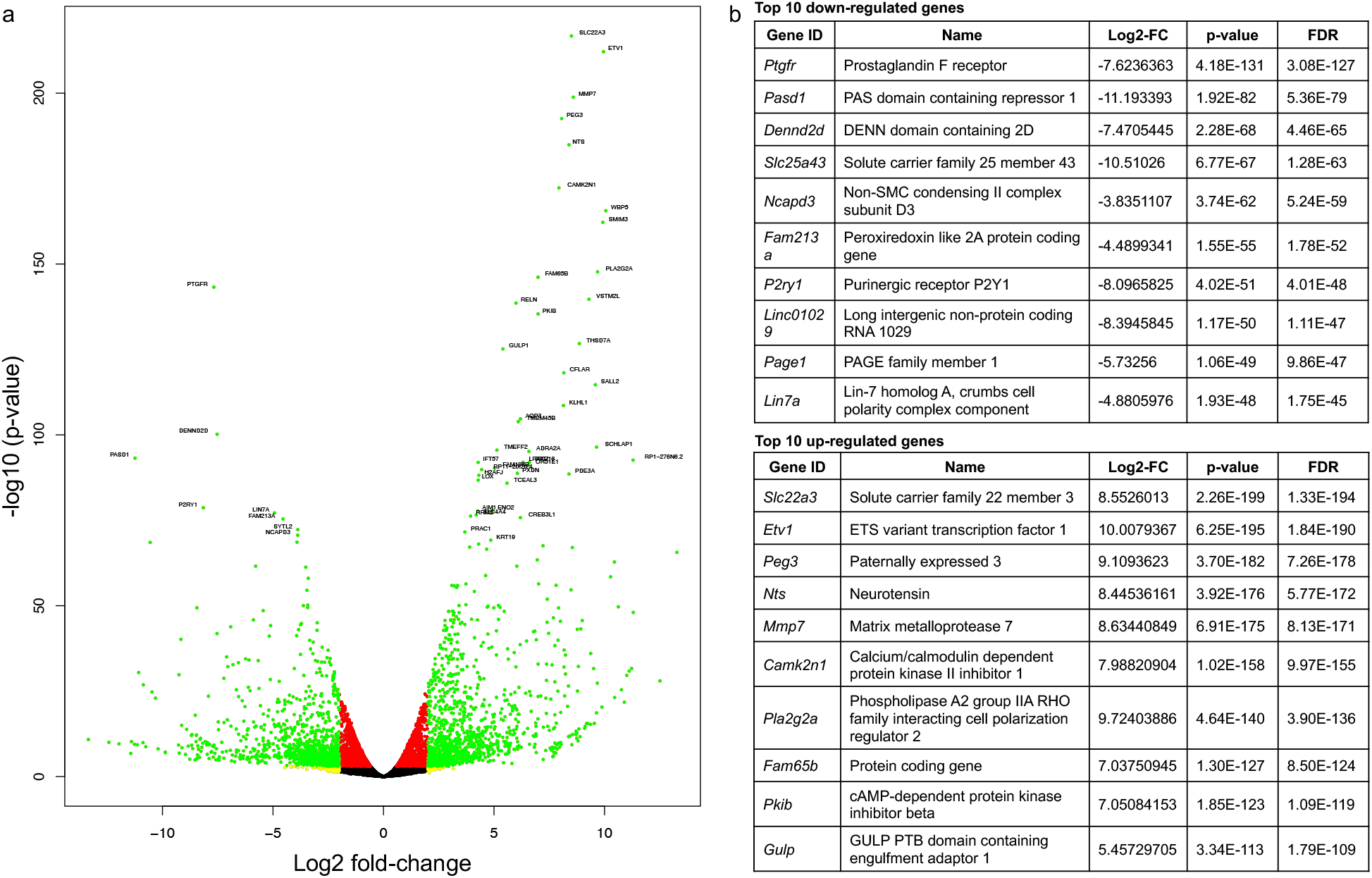
EBRT-induced transcriptomic changes in LNCaP-AR xenografts. (a) Volcano plot showing 4,851 (8.24%) DEGs (FDR=0.01) following EBRT. (b) Top 10 up- and downregulated genes (FDR=0.01).

### EBRT increases AR activity in PCa *in vivo*

To confirm EBRT-induced AR signaling *in vivo*, activation of an AR-reporter gene in LNCaP-AR tumors was assessed using bioluminescence imaging. EBRT increased mean AR-activity without significant differences between 1 and 4 fractions (**Fig. 1D**).

### [^89^Zr]11B6-uptake is an indicator of EBRT-induced AR activity

[^89^Zr]11B6 tissue uptake was assessed in 22Rv1 and LNCaP-AR tumors treated with 2, 5 or 10 Gy (1 or 4 fractions) EBRT or left untreated (**Fig. 3**). A total EBRT dose >10 Gy significantly increased uptake of [^89^Zr]11B6 by LNCaP-AR tumors (38.61-47.24 %IA/g vs. 17.9%-28.3 %IA/g in NT) and 22Rv1 xenografts (13.2-62.6 %IA/g, vs. 7.9-11.2 %IA/g NT). Co-injection of cold 11B6 significantly decreased [^89^Zr]11B6-uptake by 22Rv1 tumors after 20Gy EBRT (13.2-21.9 %IA/g vs 2.1-13.2 %IA/g blocked), confirming hK2 specificity (**Fig. 3B**).

**Figure 3.**
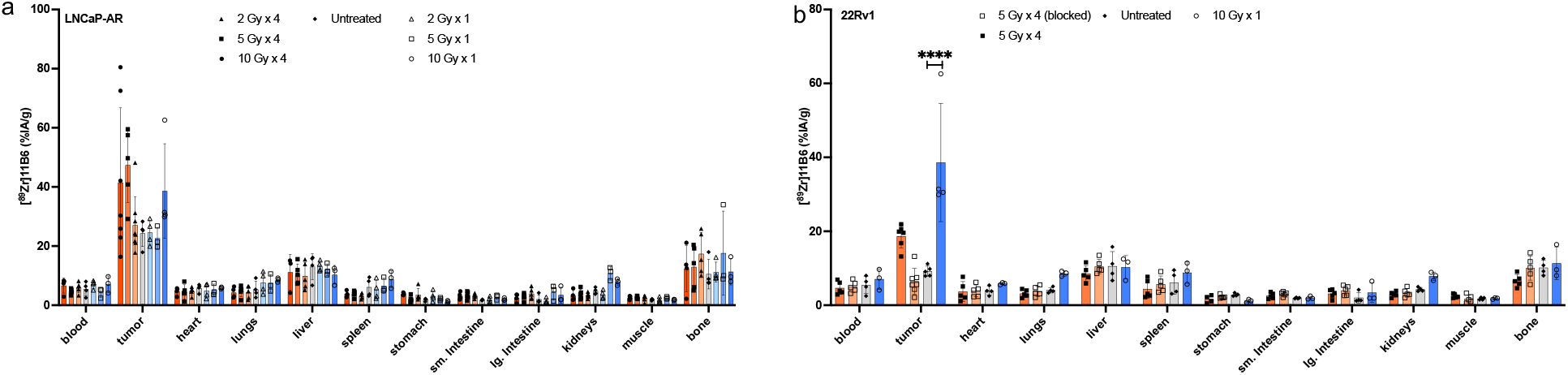
[^89^Zr]11B6 localizes to PCa after irradiation. *Ex vivo* biodistribution of [^89^Zr]11B6 in LNCaP-AR (a) and 22Rv1 (b) at 120h post-EBRT revealed higher uptake in irradiated tumors that received more than 8 Gy total dose of EBRT. Cold, unlabeled 11B6 confirmed specificity in 22Rv1. Mean ± SD and individual values are given; statistical significance was calculated for tumor uptake (NT vs. EBRT) using one-way ANOVA and Dunnett’s test for multiple comparisons.

### EBRT-induced AR activity in PCa can be monitored by [^89^Zr]11B6 positron emission tomography (PET) / computed tomography (CT) imaging

To confirm [^89^Zr]11B6 uptake as a surrogate marker for EBRT-induced AR activity, [^89^Zr]11B6 uptake was quantified *in vivo* and *ex vivo* in Pb_*KLK2* (non-malignant) and Hi-*Myc* x Pb_*KLK2* (PCa) mice after treatment with 5 fractions of 10 Gy. No significant volumetric changes were observed by MRI (**Fig. 4A**,**B**) after EBRT treatment of PCa tissue. EBRT increased AR expression in PCa (Hi-*Myc* x Pb-*KLK2*) (**Fig. 4C**); this was paralleled by significantly higher [^89^Zr]11B6 uptake after EBRT *in vivo* (%IA/g = 11.04 ± 4.42 vs. 20.23 ± 4.28).

**Figure 4.**
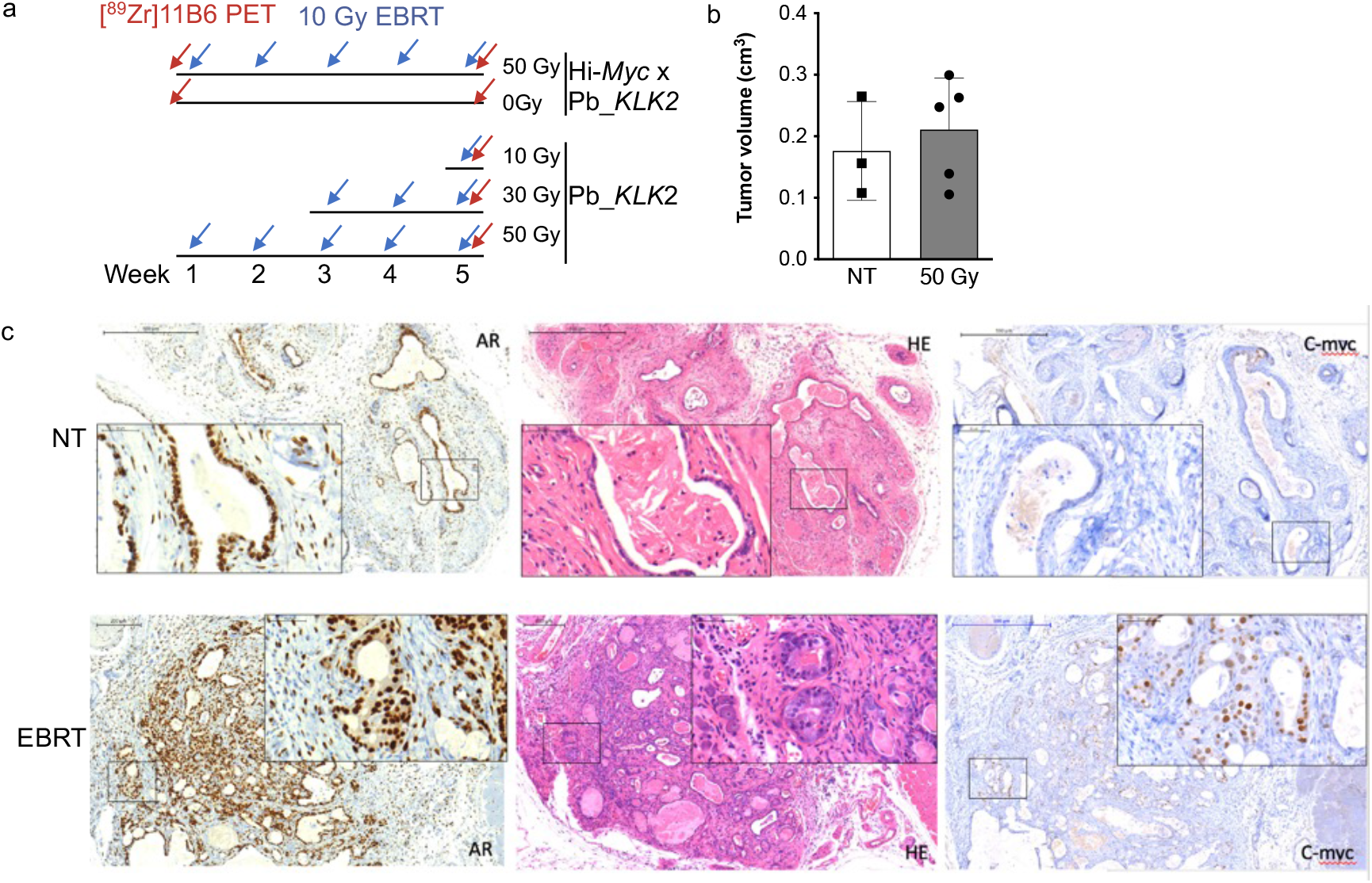
EBRT treatment of Hi-*Myc* x Pb_*KLK2* and Pb_*KLK*2 mice. (a) EBRT and imaging schedule for PCa (Hi-*Myc* x Pb_*KLK2*) and healthy (Pb_*KLK2*) mice. (b) MR imaging revealed comparable PCa volumes ±50 Gy treatment. Mean ± SD and individual values are given; statistical significance was calculated using unpaired two-tailed t-test (p=0.5872). (c) IHC of Hi-Myc x Pb-*KLK2* tumors revealed increased intratumor AR and c-MYC expression after EBRT (magnification: overview 10x, insert 40x).

In contrast, EBRT did not impact uptake in Pb_*KLK2* mice (**Fig. 5A-C**). Correlation of hK2 protein levels in tumors and [^89^Zr]11B6-uptake further confirmed AR activity (**Fig. 5D**). Taken together, these results indicate that hK2-targeted [^89^Zr]11B6 can noninvasively monitor increased AR signaling after radiotherapy in a *Myc*-driven model of PCa.

**Figure 5.**
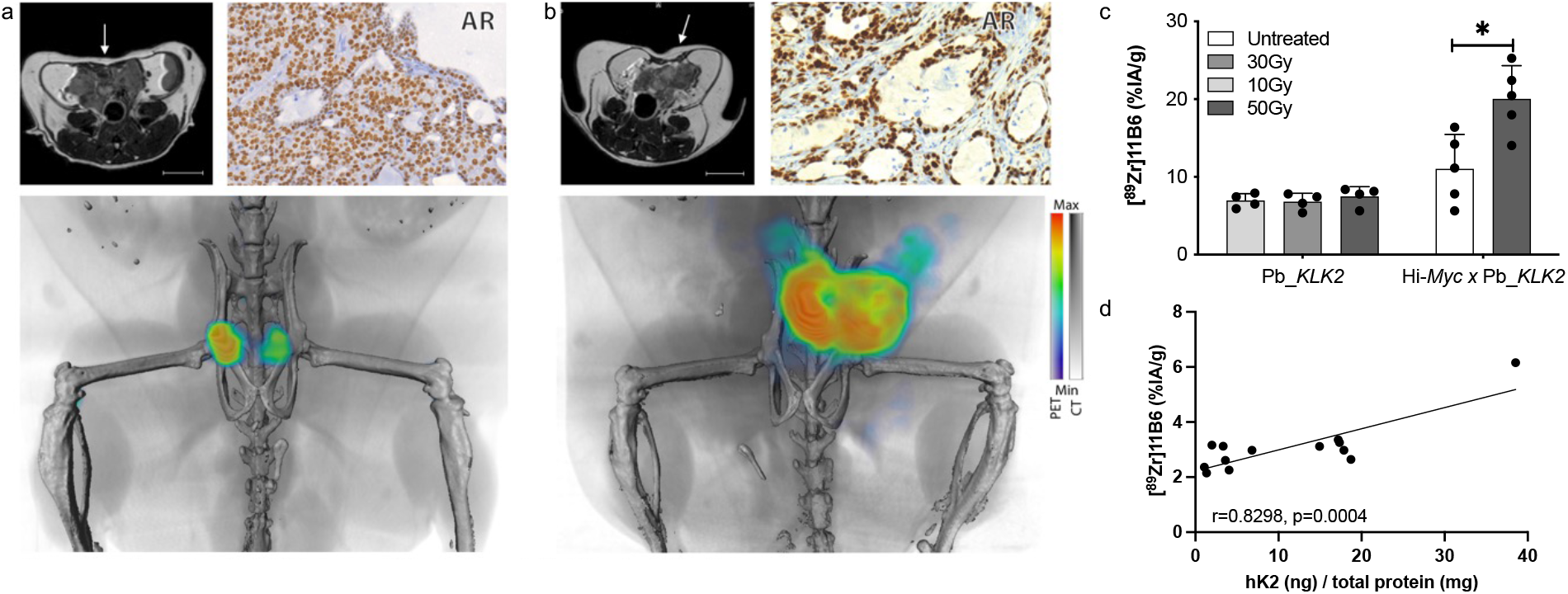
AR activity increase following EBRT visualized by [^89^Zr]11B6-PET/CT. Representative MR, IHC (40x magnification) and volume rendered PET/CT images of NT (a) and (b) 50 Gy treated Hi-*Myc* x Pb_*KLK2*. White arrow indicates prostate location in MR images (scale: 0.5 cm). (c) Activity concentration of [^89^Zr]11B6 increased following irradiation (p<0.05). Mean ± SD are given; statistical significance was calculated using unpaired two-tailed t-test. (d) PET signal from [^89^Zr]11B6 corresponds with *ex vivo* hK2 expression.

## Discussion

The current study demonstrates that EBRT-induced AR-activity, which increases in a dose-dependent manner, can be monitored noninvasively using PET. Activation of AR-signaling by EBRT may serve as prognostic biomarker and improve development of EBRT combination regimens. In a phase 3 clinical trial, the combination of EBRT with bicalutamide increased disease-free survival^24^, and PSA decay rate during salvage radiotherapy has been identified as a predictor of progression-free survival^25^. EBRT-induced AR-activity might thus negatively impact patient outcomes, and vice versa, inhibition of this response may improve patient care. Attempts to monitor AR noninvasively have been made with [^18^F]FDHT, a radio-analog of testosterone^26^; however, [^18^F]FDHT reports AR levels rather than its functional signaling activity. To measure AR pathway activity, several AR target genes are utilized as biomarkers and therapeutic targets in PCa, including prostate-specific membrane antigen (PSMA) and PSA. Recently, FDA-approved PSMA-PET has increased the ability to detect metastatic PCa lesions and is considered as a strategy to monitor AR blockade by ADT. Unfortunately, preclinical and clinical studies demonstrated that PSMA-PET is not an optimal tool for assessment of ADT efficacy^27–30^. We observed similar findings in our evaluation of PSMA levels after EBRT; *FOLH1* expression increased 2.5-fold in 22Rv1 but not in LNCaP-AR xenografts. Additionally, discrepancies in [^68^Ga]PSMA-11 PET/CT and biochemical response (PSA) limit the utility of PSMA-PET for evaluating therapeutic outcomes^30^. Taken together, these results underline the complex links between AR-activity, resistance, and AR pathway biomarkers.

*KLK2* expression and corresponding hK2 protein levels are well-established as biomarkers of AR pathway activity^12,17^. In line with a previous study^12^, we showed that EBRT increases *KLK2* expression in a dose-dependent manner. To noninvasively target *KLK2* expressing cells, we developed 11B6, an antibody that specifically internalizes into PCa cells in response to AR-activity by binding uncomplexed hK2^17^. 11B6 can be exploited for PET, single photon emission tomography, intra-operative imaging^17,31^, and radioimmunotherapy^18,26,40-41^. Studies in multiple rodent models and non-human primates showed that [^89^Zr]11B6 rapidly accumulates in PCa^18^, and changes in PCa [^89^Zr]11B6-uptake correspond to both AR-activity and hK2 protein levels^17^. We thus hypothesized that [^89^Zr]11B6 could be used to monitor changes in AR-activity during and after EBRT. We confirmed relevance of [^89^Zr]11B6-uptake as biomarker by correlating its tumor-uptake with EBRT-induced expression of the canonical AR biomarker *KLK2*. Furthermore, EBRT did not increase [^89^Zr]11B6 prostate uptake in healthy Pb_*KLK2* mice while uptake was significantly elevated in PCa of Hi-*Myc* x Pb_*KLK2* mice; this suggests that EBRT-induced AR activation is a radiobiological response unique to malignant prostate tissues.

EBRT-induced AR activation exclusively in PCa-bearing mice as well as elevated *MYC* levels in xenografts and c-MYC expression in the genetic PCa model after EBRT support the known relationship between MYC and AR. MYC upregulation has been shown to antagonize AR signaling and AR target gene expression in patient samples^32^ but has been positively correlated to AR variant expression in another study^33^. Upregulation of MYC may provide rationale for the use of co-treatment concepts using direct or indirect MYC inhibitors to block additional pro-tumorigenic transcription factors that drive PCa^34^.

The difference in [^89^Zr]11B6 uptake in the LNCaP-AR xenograft tumor model and the well-documented role of AR as a transcription factor led us to hypothesize that there would be a significant transcriptomic impact in the post EBRT-treatment setting. However, analysis of RNA-sequencing of irradiated mice revealed a downregulation of AR, highlighting the variability in tissue response to EBRT. This result exemplifies the need for diagnostic agents that focus on assessing functional AR pathway activity rather than the number of available receptors or AR expression itself. Upregulation of AR pathway target genes *KLK2* and *KLK3* in our data clearly demonstrate that the AR pathway is being differentially activated in tumor-bearing mice after radiotherapy.

The transcriptional EBRT-signature observed in the current study is in line with that reported for 11B6 alpha-radioimmunotherapy in Hi-*Myc* x Pb_*KLK2* mice^35^. Comparison of the top ten up- and downregulated DEGs revealed five common up- (*MMP7, ETV1, NTS, PLA2G2A, PEG3*) and down-regulated DEGs (*PASD1, DENN2D, PTGFR, SLC25A43, FAM213A*); this similarity underscores the ability of [^89^Zr]11B6-PET to reflect AR-driven therapeutic responses.

Overall, we demonstrated a highly specific and sensitive approach for noninvasive monitoring of functional AR-activity during and after EBRT. Exclusively in cancerous tissue, [^89^Zr]11B6 tumor-uptake correlated with AR pathway activation after irradiation. Changes in [^89^Zr]11B6 PCa-uptake paralleled increases in *KLK2* and *AR* expression seen in qPCR analysis, as well as *ex vivo* hK2 protein concentrations and IHC staining. Monitoring the AR-target gene hK2 in the treatment setting could allow patient stratification based on AR-pathway response, refinement of treatment and dosing strategies, e.g., by selection of AR-targeted treatment combinations, and may provide mechanistic insights into enhancement of EBRT in some patients with concurrent or adjuvant ARSI.

## Methods

### Radiochemistry

Radiosynthesis of [^89^Zr]-DFO-11B6 ([^89^Zr]11B6) has previously been described^31^. 11B6 antibody was provided by Dr. Kim Pettersson, University of Turku, Finland. All labeling reactions achieved >99% radiochemical purity. Average specific activity of the final radiolabeled conjugate was 51.8 MBq/mg (1.4 mCi/mg).

### Cell lines

22Rv1 cells were purchased from ATCC. LNCaP-AR (LNCaP with overexpression of wildtype AR) was a kind gift from Charles Sawyers^36^. Cells were cultured according to the providers’ instructions and frequently tested for mycoplasma contamination.

### Mouse models

All animal experiments were conducted in compliance with MSKCC guidelines, IACUC-established guidelines, and RARC animal protocol (# 04-01-002). Xenografts were established in male athymic BALB/c (nu/nu) mice (6–8 weeks old, 20-25 g; Charles River) by subcutaneous injection of LNCaP-AR or 22Rv1 cells (1-5×10^6^ cells, 1:1 = media : Matrigel). Tumors developed after 3-7 weeks. The transgenic PCa mouse models used, Hi-*Myc* × Pb_*KLK2* with prostate-specific AR-driven hK2 expression, as well as Pb_*KLK2* mice with abundant AR-driven hK2 expression specific to murine prostate tissue, have been previously reported^17^.

### EBRT

Irradiation of disease sites was performed as previously described^37^. Briefly, a whole-body CT was acquired (XRad225Cx, Precision X-Ray, Inc.; dual focal spot x-ray tube at 45 kVp with a flat-panel amorphous silicon imager mounted on a C-arm gantry), tumor fields were identified and a treatment plan with >3 angles and a dose rate of ∼3 Gy/min (tube voltage, 225 kVp) was devised. Radiation dosimetry was performed using Gafchromic EBT film (ISP Inc.); a clear film that polymerizes with increasing optical density to a degree linearly with dose. The Gafchromic film verified the targeting accuracy, the magnitude of dose delivered and the geometry of the planned dose plan.

### Magnetic resonance imaging

Prostate tumor volumes were defined using T2-weighted MR scans (Bruker BioSpin 4.7 T). An interleaved T2-weighted turbo spin echo sequence (3,200/57.1) with 8 averages was used, with slice dimensions of 8.5 × 3.99 × 0.8 cm. A total scan duration of 10 minutes 14 seconds generated 220 µm and 800 μm in and out of plane slices, respectively. A trained reader calculated prostate volumes by segmenting the prostate (OsiriX, v8.1)^38^.

### Gene expression analysis

RNA was purified using the RNeasy Mini Kit (Qiagen), and quantitative PCR to determine expression of *KLK2, KLK3*, and *FOLH1* was performed as previously described.

For RNA-sequencing, raw read count RNA-sequencing data were generated from untreated (NT; n = 3) LNCaP-AR tumor samples and 5 × 10 Gy (n=3) treated samples. A total of 58,828 genes were acquired and analyzed as previously reported^35^. Both hierarchical clustering analysis (based on Euclidean distance) and multi-dimensional scaling (MDS) plots demonstrated a clear division between the samples from the two cohorts (**Suppl. Fig. 1, 2**). Differentially expressed genes (DEGs) were defined at an adjusted p<0.001 and an absolute value of log2 fold-change >1. A positive fold-change represented up- and a negative fold change represented downregulation in EBRT-treated tumors. Pathway analysis was performed using enrichR^39^ and the KEGG 2021 database.

### Bioluminescence imaging

Activity of the AR-dependent reporter construct expressed in LNCaP-AR tumors was quantified by bioluminescence imaging (Living Image^®^ 4.5.2) following retro-orbital injection of D-Luciferin (30 mg/mL, 10 µL; exposure times 1, 5, 10, 20, and 40 seconds). Data were expressed as radiance (photons/s) divided by tumor volume measured by caliper (V = length x width^2^).

### Impact of EBRT on [^89^Zr]11B6 tumor uptake

Mice bearing LNCaP-AR and 22Rv1 xenografts, and Hi-*Myc* × Pb_*KLK2* and Pb_*KLK2* mice, received [^89^Zr]11B6 (3.7–5.55 MBq [100–150 μCi], 25 µg protein, i.v.; t=0 h), after EBRT (n=4– 5/group). To confirm specificity, a control group of mice with 22Rv1 tumors treated with 4 × 5 Gy was co-injected with 1 mg of unlabeled 11B6. [^89^Zr] radioactivity in tumors and organs harvested 120 h post-injection (p.i.) was quantified using a gamma-counter. Data were background and decay corrected, and the percentage injected activity per gram tissue (%IA/g) was calculated.

### Monitoring AR-activity using PET/ CT

PET/CT imaging (Inveon MM, IRW Acquisition software) was performed as previously described^40^, at 120 h p.i. with Hi-*Myc* × Pb_*KLK*2 following administration of [^89^Zr]11B6 (3.7–5.55 MBq [100–150 μCi], 25 µg of protein, i.v.). Duration of PET scans were ∼1 h or until 20 × 10^6^ coincident events were recorded. A 3D maximum a priori reconstruction was used to generate tomographic datasets. Assessment of hK2 expression for correlation with [^89^Zr]11B6 uptake was reported previously^17^.

### Histology

Prostate tissues of Hi-*Myc* × Pb_*KLK2* and Pb_*KLK2* mice harvested after EBRT (5 × 10 Gy) were fixed in 4% paraformaldehyde and cut into 15 µm sections before staining with hematoxylin and eosin (H&E). Immunohistochemistry (IHC) for detection of AR and c-MYC was performed at the Molecular Cytology Core Facility (MSKCC) using a Discovery XT processor (Ventana Medical Systems). Sections were blocked in 10% normal goat serum in PBS for 30 minutes before staining with an anti-AR (N-20) antibody (1 µg/mL, 3 h; Santa Cruz, #SC-816; secondary: biotinylated goat anti-rabbit IgG, 1:200, 16 minutes; Vector labs, #PK6101), or an anti-c-MYC antibody (1:100, 5h; Epitomics, #P01106; secondary: biotinylated goat anti-rabbit IgG, 1:200, 1 h; Vector labs, #PK6101). Blocker D, Streptavidin-HRP and DAB detection kit (Ventana Medical Systems) were used according to the manufacturer’s instructions.

### Statistics

Statistical significance was determined by unpaired two-tailed t-test (2 groups) or, for >2 groups, by one-way ANOVA followed by Dunnett’s test to correct for multiple comparisons and set to p<0.05. Data are presented as mean ± standard deviation (SD). Analysis was performed with GraphPad Prism Version 9.2.0. For RNA-sequencing, differentially expressed genes were considered significant with an adjusted *p*<0.001 and log2 fold-change >1 as described previously^35^.

## Acknowledgements

We thank Drs. Brent Rupnow, Marco Gottardis, and Michael Russell at oncology innovation at the Janssen pharmaceutical companies of Johnson & Johnson for excellent discussions and intellectual input.

This study was supported in part by Janssen R&D LLC, UCLA Eli and Edythe Broad Center of Regenerative Medicine and Stem Cell Research Rose Hill Foundation Innovator Award, UCLA SPORE in prostate cancer (P50 CA092131), the Imaging and Radiation Sciences Program, U.S. NIH grant P30 CA008748 (MSKCC Support Grant). The MSKCC Small-Animal Imaging Core Facility is supported in part by NIH grants P30 CA008748-48, S10 RR020892-01, S10 RR028889-01, and the Geoffrey Beene Cancer Research Center. We also acknowledge William H. Goodwin and Alice Goodwin and the Commonwealth Foundation for Cancer Research, the Experimental Therapeutics Center, and the Radiochemistry & Molecular Imaging Probe Core (P50-CA086438), all of MSKCC. M.R. McDevitt: NIH R01CA166078, R01CA55349, P30CA008748, P01CA33049, F31CA167863, the MSKCC for Molecular Imaging and Nanotechnology. D.L.J. Thorek: NCI R01CA201035, R01CA240711, and R01CA229893. H. Lilja: NIH/NCI CCSG to MSKCC (P30 CA008748). S.M. Larson: Ludwig Center for Cancer Immunotherapy (MSKCC), NCI P50-CA86438. H. Lilja: NIH/NCI P50CA092629 and R01 CA244948 Sidney Kimmel Center for Prostate and Urologic Cancers. David H. Koch Prostate Cancer Foundation Award, Swedish Cancer Society (Cancerfonden 20 1354 PjF), Swedish Research Council (VR-MH 2016-02974), General Hospital in Malmö Foundation for Combating Cancer. D.L.J. Thorek: NCI R01CA201035, R01CA240711, R01CA229893. D. Ulmert, M.R. McDevitt: DoD W81XWH-18-1-0223. D. Ulmert: UCLA SPORE in Prostate Cancer (P50 CA092131), JCCC Cancer support grant from NIH P30 CA016042 (PI: Teitell), Knut and Alice Wallenberg Foundation, Bertha Kamprad Foundation, David H. Koch Prostate Cancer Foundation Young Investigator Award, Swedish Cancer Society, and Swedish Cancer Foundation.

## Author contributions

D.L.J.T. and D.U. conceived, designed, performed, and analyzed all the experiments and data. C.M.S, M.A., M.B. D.V. and K.L. performed the data acquisition and analysis; D.R.V., M.R.M. and D.A. assisted in study design and radioconjugate formulation. T.K., J.E.P., K.H. W.Z., N.P., R.J.K assisted in technical study design and data analysis. R.D., S.M.L. and H.S. supervised the aspects of the project. All authors discussed the results, prepared, commented and approved the manuscript.

## References

1. Chen, Y., Sawyers, C. L. & Scher, H. I. Targeting the androgen receptor pathway in prostate cancer. Curr. Opin. Pharmacol. 8, 440–448 (2008).

2. Huggins, C. & Hodges, C. V. Studies on prostatic cancer: I. The effect of castration, of estrogen and of androgen injection on serum phosphatases in metastatic carcinoma of the prostate. 1941. J. Urol. 168, 9–12 (2002).

3. Moris, L. et al. Benefits and Risks of Primary Treatments for High-risk Localized and Locally Advanced Prostate Cancer: An International Multidisciplinary Systematic Review. Eur. Urol. 77, 614–627 (2020).

4. Bolla, M. et al. External irradiation with or without long-term androgen suppression for prostate cancer with high metastatic risk: 10-year results of an EORTC randomised study. Lancet Oncol. 11, 1066–1073 (2010).

5. Widmark, A. et al. Endocrine treatment, with or without radiotherapy, in locally advanced prostate cancer (SPCG-7/SFUO-3): an open randomised phase III trial. Lancet 373, 301– 308 (2009).

6. Polkinghorn, W. R. et al. Androgen receptor signaling regulates DNA repair in prostate cancers. Cancer Discov. 3, 1245–1253 (2013).

7. Goodwin, J. F. et al. A hormone-DNA repair circuit governs the response to genotoxic insult. Cancer Discov. 3, 1254–1271 (2013).

8. Arora, V. K. et al. Glucocorticoid receptor confers resistance to antiandrogens by bypassing androgen receptor blockade. Cell 155, 1309–1322 (2013).

9. Handle, F. et al. Drivers of AR indifferent anti-androgen resistance in prostate cancer cells. Sci. Rep. 9, 13786 (2019).

10. Antonarakis, E. S. et al. AR-V7 and resistance to enzalutamide and abiraterone in prostate cancer. N. Engl. J. Med. 371, 1028–1038 (2014).

11. Zhu, Y. et al. Role of androgen receptor splice variant-7 (AR-V7) in prostate cancer resistance to 2nd-generation androgen receptor signaling inhibitors. Oncogene 39, 6935– 6949 (2020).

12. Spratt, D. E. et al. Androgen Receptor Upregulation Mediates Radioresistance after Ionizing Radiation. Cancer Res. 75, 4688–4696 (2015).

13. Chaiswing, L., Weiss, H. L., Jayswal, R. D., Clair, D. K. S. & Kyprianou, N. Profiles of Radioresistance Mechanisms in Prostate Cancer. Crit. Rev. Oncog. 23, 39–67 (2018).

14. Ulmert, D., O’Brien, M. F., Bjartell, A. S. & Lilja, H. Prostate kallikrein markers in diagnosis, risk stratification and prognosis. Nat. Rev. Urol. 6, 384–391 (2009).

15. Prostate-specific kallikrein-related peptidases and their relation to prostate cancer biology and detection. Established relevance and emerging roles. Thromb. Haemost.

16. Sävblom, C. et al. Blood levels of free-PSA but not complex-PSA significantly correlates to prostate release of PSA in semen in young men, while blood levels of complex-PSA, but not free-PSA increase with age. Prostate 65, 66–72 (2005).

17. Thorek, D. L. J. et al. Internalization of secreted antigen-targeted antibodies by the neonatal Fc receptor for precision imaging of the androgen receptor axis. Sci. Transl. Med. 8, 367ra167 (11 30, 2016).

18. Feed-forward alpha particle radiotherapy ablates androgen receptor-addicted prostate cancer. Nat. Commun.

19. Thorek, D. L. J. et al. Harnessing Androgen Receptor Pathway Activation for Targeted Alpha Particle Radioimmunotherapy of Breast Cancer. Clin. Cancer Res. 25, 881–891 (2019).

20. Baena, E. et al. ETV1 directs androgen metabolism and confers aggressive prostate cancer in targeted mice and patients. Genes Dev. 27, 683–698 (2013).

21. Sharma, N. L. et al. The androgen receptor induces a distinct transcriptional program in castration-resistant prostate cancer in man. Cancer Cell 23, 35–47 (2013).

22. Litvinov, I. V. et al. Androgen receptor as a licensing factor for DNA replication in androgen-sensitive prostate cancer cells. Proc. Natl. Acad. Sci. U. S. A. 103, 15085–15090 (2006).

23. Schiewer, M. J. et al. Dual roles of PARP-1 promote cancer growth and progression. Cancer Discov. 2, 1134–1149 (2012).

24. Radiation Therapy With or Without Bicalutamide and Goserelin in Treating Patients With Prostate Cancer. ClinicalTrials.gov identifier: NCT00021450 (2016).

25. Gunnlaugsson, A. et al. PSA decay during salvage radiotherapy for prostate cancer as a predictor of disease outcome - 5 year follow-up of a prospective observational study. Clin Transl Radiat Oncol 24, 23–28 (2020).

26. Beattie, B. J. et al. Pharmacokinetic assessment of the uptake of 16beta-18F-fluoro-5alpha-dihydrotestosterone (FDHT) in prostate tumors as measured by PET. J. Nucl. Med. 51, 183–192 (2010).

27. Enzalutamide With Lu PSMA-617 Versus Enzalutamide Alone in Men With Metastatic Castration-resistant Prostate Cancer (ENZA-p). ClinicalTrials.gov Identifier: NCT04419402 (2022).

28. Lückerath, K. et al. Preclinical evaluation of PSMA expression in response to androgen receptor blockade for theranostics in prostate cancer. EJNMMI Res. 8, 96 (2018).

29. Staniszewska, M. et al. Enzalutamide Enhances PSMA Expression of PSMA-Low Prostate Cancer. Int. J. Mol. Sci. 22, 7431 (2021).

30. Kuten, J., Sarid, D., Yossepowitch, O., Mabjeesh, N. J. & Even-Sapir, E. [68Ga]Ga-PSMA-11 PET/CT for monitoring response to treatment in metastatic prostate cancer: is there any added value over standard follow-up? EJNMMI Res. 9, 84 (2019).

31. Timmermand, O. V. et al. Preclinical imaging of kallikrein-related peptidase 2 (hK2) in prostate cancer with a (111)In-radiolabelled monoclonal antibody, 11B6. EJNMMI Res. 4, 51 (2014).

32. Barfeld, S. J. et al. c-Myc Antagonises the Transcriptional Activity of the Androgen Receptor in Prostate Cancer Affecting Key Gene Networks. EBioMedicine 18, 83–93 (2017).

33. Bai, S. et al. A positive role of c-Myc in regulating androgen receptor and its splice variants in prostate cancer. Oncogene 38, 4977–4989 (2019).

34. Llombart, V. & Mansour, M. R. Therapeutic targeting of “undruggable” MYC. EBioMedicine 75, 103756 (2022).

35. Bicak, M. et al. Genetic signature of prostate cancer mouse models resistant to optimized hK2 targeted α-particle therapy. Proc. Natl. Acad. Sci. U. S. A. 117, 15172–15181 (2020).

36. Chen, C. D. et al. Molecular determinants of resistance to antiandrogen therapy. Nat. Med. 10, 33–39 (2004).

37. Thorek, D. L. J. et al. Reverse-Contrast Imaging and Targeted Radiation Therapy of Advanced Pancreatic Cancer Models. Int. J. Radiat. Oncol. Biol. Phys. 93, 444–453 (2015).

38. Rosset, A., Spadola, L. & Ratib, O. OsiriX: an open-source software for navigating in multidimensional DICOM images. J. Digit. Imaging 17, 205–216 (2004).

39. Kuleshov, M. V. et al. Enrichr: a comprehensive gene set enrichment analysis web server 2016 update. Nucleic Acids Res. 44, W90–7 (2016).

40. Veach, D. R. et al. PSA-Targeted Alpha-, Beta-, and Positron-Emitting Immunotheranostics in Murine Prostate Cancer Models and Nonhuman Primates. Clin. Cancer Res. (2021) doi:10.1158/1078-0432.CCR-20-3614.

