## Supplementary Figures for "Assessing Functional Androgen Receptor Pathway Activity in Response to Radiotherapy Using hK2-targeted PET Imaging"

- **Supplementary Data**

Running title: hK2-PET for EBRT assessment

Claire M Storey[1], Mohamed Altai[2], Mesude Bicak[3], Darren R Veach[4], Katharina Lückerath[5], Gabriel Adrian[2], Michael R McDevitt[4], Teja Kalidindi[4], Julie E Park[1], Ken Herrmann[5], Diane Abou[6,7], Wahed Zedan[2], Norbert Peekhaus[2], Robert J Klein[8], Robert Damoiseaux[1,9], Steven M Larson[4,10], Hans Lilja[11-14], Daniel Thorek[6,7,15], David Ulmert*[1,2,9,16-18]

[1] Department of Molecular & Medical Pharmacology, University of California Los Angeles (UCLA), Los Angeles, USA

[2] Division of Oncology and Pathology, Department of Clinical Sciences, Lund University, Lund, Sweden

[3] Hasso Plattner Institute for Digital Health, Department of Genetics and Genomic Sciences, Icahn School of Medicine at Mount Sinai, New York, USA

[4] Department of Radiology, Memorial Sloan Kettering Cancer Center (MSKCC), New York, USA

[5] Department of Nuclear Medicine, University Hospital Essen, University of Duisburg-Essen, DKTK, Essen, Germany

[6] Department of Radiology, Washington University School of Medicine, St. Louis, USA

[7] Siteman Cancer Center, Washington University School of Medicine, St. Louis, USA

[8] Icahn Institute for Genomics and Multiscale Biology, Department of Genetics and Genomic Sciences, Icahn School of Medicine at Mount Sinai, New York, USA

[9] California NanoSystems Institute, UCLA, Los Angeles, USA

[10] Department of Radiology, Weill Cornell Medical College, New York, USA

[11] Genitourinary Oncology Service, Department of Medicine, MSKCC, New York, USA

[12] Urology Service, Department of Surgery, MSKCC, New York, USA

[13] Department of Laboratory Medicine, MSKCC, New York, NY 10065, USA

[14] Department of Translational Medicine, Lund University, Malmö, Sweden

[15] Department of Biomedical Engineering, Washington University School of Medicine, St. Louis, USA

[16] Department of Urology, Institute of Urologic Oncology, UCLA, Los Angeles, USA

[17] Jonsson Comprehensive Cancer Center, David Geffen School of Medicine, UCLA, Los Angeles, USA

[18] Eli and Edythe Broad Center of Regenerative Medicine and Stem Cell Research, UCLA, Los Angeles, USA

Corresponding author:

David Ulmert

650 Charles E Young Drive S, Los Angeles, CA 90095, USA

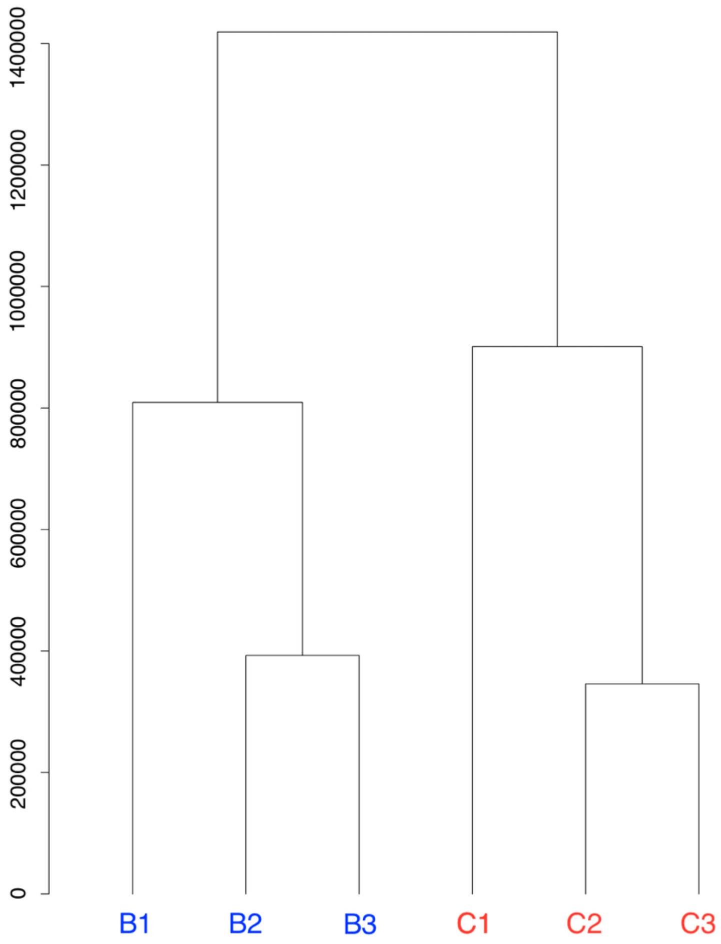


**Suppl. Fig 1. Hierarchical clustering based on Euclidean distance of RNA-seq data demonstrates separation of EBRT-treated (B1-3) and untreated (C1-3) samples.**

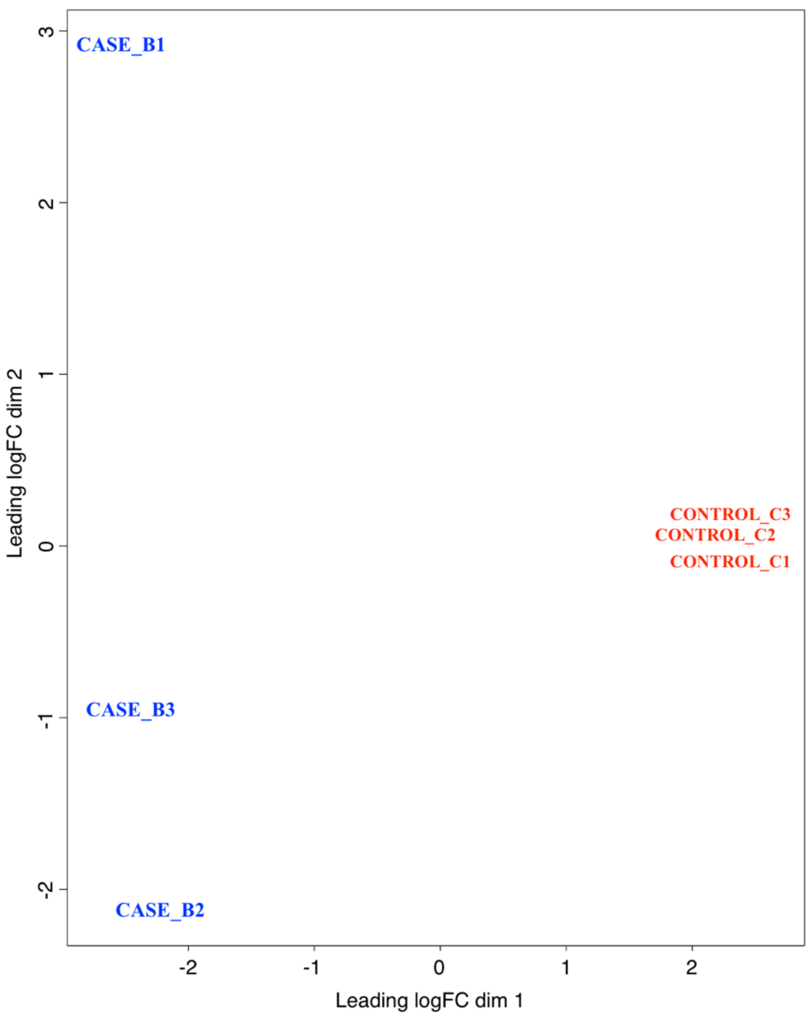


**Suppl. Fig 2. Multi-dimensional scaling plot of RNA-seq data demonstrates separation of EBRT-treated (B1-3) and untreated (C1-3) samples.**
